# STRUCTURAL PLASTICITY ALLOWS DOUBLECORTIN AND DRUG-STABILIZED MICROTUBULES TO ACCOMMODATE EACH OTHER

**DOI:** 10.64898/2026.11.03.763982

**Authors:** Varun Ramaswamy, Szymon W. Manka, Darsh D. Sharma, Anna Akhmanova, Ankit Rai, Carolyn A. Moores

## Abstract

Microtubule architecture and dynamics are modulated by small molecules, but how these compounds alter lattice properties and interactions with cellular regulators remains incompletely understood. We used the neuronal microtubule-associated protein doublecortin (DCX), previously reported to preferentially recognize 13-protofilament microtubules, to probe lattices stabilized by different taxanes, including the fluorescent taxane Fchitax-3. We show that DCX binds 15-protofilament Fchitax-3-stabilized microtubules, demonstrating that DCX is not a specific marker of 13-protofilament architecture. Cryo-electron microscopy revealed that the fluorescent moiety of Fchitax-3 inserts into inter-protofilament lateral contacts, explaining its effects on microtubule architecture and dynamics. DCX binds Fchitax-3 microtubules through its N-terminal DC domain, which accommodates the narrower inter-protofilament binding site in the 15-protofilament lattice. Remarkably, although Fchitax-3 induces the longitudinal lattice expansion characteristic of taxanes, DCX remodels its binding site towards its preferred compacted configuration. Thus, taxanes can modify microtubule structure in distinct ways while preserving lattice plasticity and the capacity to undergo conformational remodelling by microtubule-associated proteins. This contrasts with the comparatively rigid lattice of GMPCPP-stabilized microtubules. Consequently, interpretations of microtubule lattice states based on comparisons between taxane- and GMPCPP-stabilized microtubules may conflate structural differences with different degrees of lattice plasticity. Our findings establish the microtubule lattice as a malleable structural substrate whose drug-induced conformations can be further remodelled by cellular binding partners.

## INTRODUCTION

Microtubules are dynamic cytoskeleton components involved in diverse processes, including trafficking and cell division. Their dynamics are driven by the *a*β-tubulin GTPase cycle that couples nucleotide state to tubulin conformation and lattice stability. GTP-bound β-tubulin stimulates nucleation and polymerization to form the hollow microtubule wall, whereas the gradual hydrolysis of GTP bound to β-tubulin promotes weakening of polymer contacts until catastrophe causes depolymerization^1^. Microtubule functions are disrupted in numerous diseases, and they are also important therapeutic targets^2,3^. Drugs that target microtubules can be broadly classified as microtubule-stabilizing agents (MSAs, e.g. Taxol®) and microtubule-destabilizing agents (MDAs, e.g. Velban®)^3^. Paclitaxel and its derivatives (known as taxanes) are widely used MSAs that bind a hydrophobic pocket on the lumenal surface of β-tubulin^4^. This binding pocket is partially occluded in unpolymerized tubulin but becomes accessible in lattice-compatible, polymerized tubulin^5^. By preferentially binding the microtubule lattice, taxanes suppress depolymerization and stabilize the polymer^6^. Beyond stabilization, the effects of taxanes on microtubule structure and dynamics have been extensively – although not always consistently – characterized. Taxanes can influence microtubule architecture, lattice expansion/compaction, polymer continuity, and long-range dynamics^5,7–9^. Exploring the interplay between taxanes and microtubule regulatory proteins may clarify the molecular mechanisms underlying the diversity of reported taxane action.

Doublecortin (DCX) is an essential neuronal MAP that nucleates and stabilizes microtubules and is required for neuronal migration during brain development^10,11^. DCX is composed of a tandem of conserved, microtubule-binding DC domains, N-DC and C-DC^12,13^. Both N-DC and C-DC can bind at the corner of four tubulin dimers on the outer surface of the microtubule lattice, bridging longitudinal and lateral lattice contacts^14,15^. DCX binding in the inter-protofilament cleft is predicted to be sensitive to microtubule protofilament number, which affects the inter-protofilament angle and therefore the space for DCX binding. Indeed, *in vitro* microtubule nucleation in the presence of DCX typically produces 13-protofilament microtubules, which is the main architecture found in cells^16^. DCX has thus been considered a marker to distinguish between 13- and non-13-protofiilament architectures^17,18^. However, numerous studies have also shown that DCX can bind to a broader range of architectures in pre-formed microtubules, including taxane-stabilized microtubules, indicating that the precise nature of its sensitivity to different microtubule substrates may be less clear cut^12,19,20^. In addition, although DCX and taxanes do not compete for the same microtubule binding site, their interplay on microtubules is complex and can be mutually exclusive^21–23^. Taken together, these observations suggest that DCX-microtubule binding in the presence of taxanes is influenced by a complex set of factors, including protofilament number, nucleotide state, exact taxane identity, and the mechanistic integration of these properties has yet to be fully resolved.

We therefore set out to explore the interplay between DCX binding and microtubules stabilized by, or seeded from, a range of taxanes. Using Total Internal Reflection Fluorescence (TIRF) microscopy, we show that DCX binds to a variety of microtubule architectures, including the 15-protofilament microtubules stabilized by the fluorescent taxane Fchitax-3. Cryo-electron microscopy (cryo-EM) structures of DCX-bound Fchitax-3-microtubules show that the fluorescent moiety of Fchitax-3 inserts at lateral contacts, thereby explaining its influence on microtubule architecture. Furthermore, our reconstructions show that while the N-DC domain of DCX flexibly adapts to the binding site found in 15-protofilament microtubules, it also moulds the microtubule lattice to facilitate binding. Together, these data reveal the unexpected adaptability of microtubule polymers and their cellular binding partners to modulation by drug-like ligands.

## RESULTS

### DCX binds a range of protofilament architectures including 15-protofilament Fchitax-3-stabilized microtubules

We first set out to test whether fluorescently tagged DCX could be used as a probe of microtubule architecture using TIRF microscopy. To this end, DCX was N-terminally tagged with TEV-bio and mCherry, purified from HEK293 cells (Fig. S1A), and added to in vitro-polymerized microtubules stabilized with different MSAs, which promote distinct microtubule conformations^9^. We also included GMPCPP-stabilized microtubules, which predominantly adopt a 14-protofilament architecture^24^ and have previously been reported to exclude DCX binding^18^. Consistent with previous work, DCX bound robustly to paclitaxel-stabilized microtubules, which in our hands predominantly contain 12 or 13 protofilaments^9^, while completely avoiding GMPCPP-stabilized microtubules (Fig. 1A, B). Surprisingly, however, DCX also associated efficiently with docetaxel-stabilized microtubules, which preferentially adopt a 14-protofilament architecture, and with Fchitax-3-stabilized microtubules, which contain 15 or 16 protofilaments^9^ (Fig. 1A, B). Notably, DCX decoration was discontinuous along individual taxane-stabilized microtubules, with alternating regions of high and low fluorescence intensity, indicating heterogeneous lattice occupancy rather than uniform binding (Fig. 1A and Fig. S1B). Since taxane-stabilized microtubules can contain structural defects, we next examined DCX binding to dynamic microtubules nucleated from GMPCPP or MSA-stabilized seeds. EB3-GFP was included to mark growing microtubule ends. Under all conditions tested, 50 nM mCherry-DCX robustly decorated the dynamic microtubule lattice, including microtubules grown from GMPCPP seeds, and displayed modest enrichment at growing plus ends, while consistently avoiding the GMPCPP-stabilized seeds (Fig. 1C). Previous work interpreted the ability of DCX to bind the dynamic lattice extending from GMPCPP seeds as evidence for a transition from a 14- to a 13-protofilament architecture at the seed boundary^18^. However, recent interference reflection microscopy (IRM) analysis did not support this interpretation, instead showing that GMPCPP-stabilized 14-protofilament seeds are extended by lattices that retain a 14-protofilament architecture^25^. Together, these findings indicate that DCX is sensitive to microtubule lattice conformation, but this sensitivity is unlikely to be determined by protofilament number.

**Figure 1.**
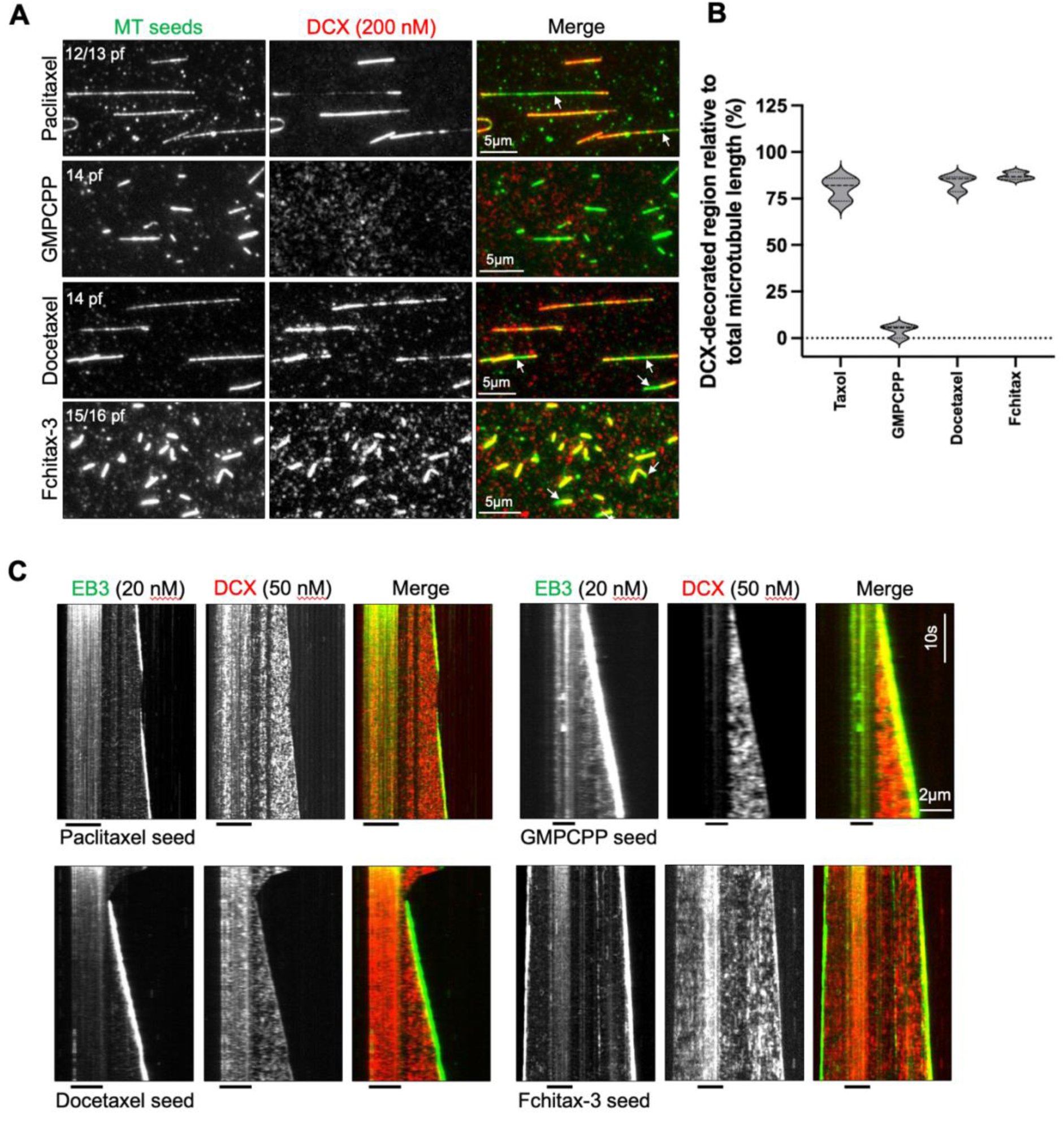
Doublecortin recognizes distinct microtubule lattice architectures independent of protofilament number. A) Representative TIRF microscopy images showing the binding of mCherry-DCX (DCX, 200 nM) to stabilized microtubule seeds assembled in the presence of paclitaxel (13-protofilament, (pf)), GMPCPP (14-pf), docetaxel (14-pf), and Fchitax-3 (15/16-pf). DCX binds to paclitaxel-, docetaxel-, and Fchitax-3-stabilized microtubules, whereas no binding is observed to GMPCPP-stabilized microtubules. White arrows in the merged images highlight regions within microtubules where DCX binding is either reduced or absent, suggesting that DCX binding is sensitive to local structural variations within the microtubule lattice. The experiment was repeated twice. B) Violin plot showing DCX lattice occupancy on microtubule seeds stabilized with paclitaxel (n = 74), GMPCPP (n = 104), docetaxel (n = 63) and Fchitax-3 (n = 137) from 3 independent sets of experiments. The respective length of microtubule lattices decorated by mCherry-DCX was measured and expressed as a percentage of the total measured microtubule length. For each dataset, the thicker dashed line denotes the median flanked by thinner dotted lines to denote first and third quartiles. C) Representative kymographs showing TIRF microscopy-based imaging of dynamic microtubules grown from paclitaxel-, GMPCPP-, docetaxel-, and Fchitax-3-stabilized seeds in the presence of DCX (50 nM) and EB3 (20 nM). DCX binds robustly to the lattices of growing microtubules regardless of the seed type. Although DCX does not associate with GMPCPP-stabilized seeds, it readily binds to the newly polymerized microtubule lattice extending from GMPCPP seeds. The position of the seed is indicated by a black line under each kymograph. Experiment was repeated twice independently.

### Perturbation of microtubule lateral contacts by the Fchitax-3 fluorophore favors higher protofilament numbers

To investigate the structural basis of the unexpectedly broad microtubule binding of DCX, we collected cryo-EM data of Fchitax-3-stabilized microtubules bound by recombinant, untagged DCX^15^. These microtubules were first characterized according to their protofilament architecture, both in terms of protofilament and start number, which defines the presence or absence of a lattice seam^26^. As observed previously, the largest proportion of the Fchitax-3-microtubules had 15 protofilaments, both with (15_3) and without (15_4) a seam^9^ (Fig. 2A). As noted above, this is distinct from the 12 and 13 protofilament architecture observed in paclitaxel-stabilized microtubules^9,27^, reinforcing the idea that modifications of taxanes can alter the architecture of the resulting stabilized polymers. Data for the 15_3 and 15_4 sub-populations were independently processed and averaged for 3D reconstruction (Fig. 2B, Fig. S2A,B, see Methods). Both reconstructions exhibited very similar overall resolutions (15_4, 3.6 Å; 15_3, 3.6 Å, Fig. S2A), and modelling of the asymmetric units showed DCX density corresponding to a characteristic single DC domain bound at the corner of four tubulin dimers making homotypic lateral contacts^14^ (Fig. 2C; see detailed analysis below). As expected, however, DCX density was absent from the heterotypic tubulin contacts at the seam of the 15_3 reconstruction^14^ (Fig. S3).

**Figure 2.**
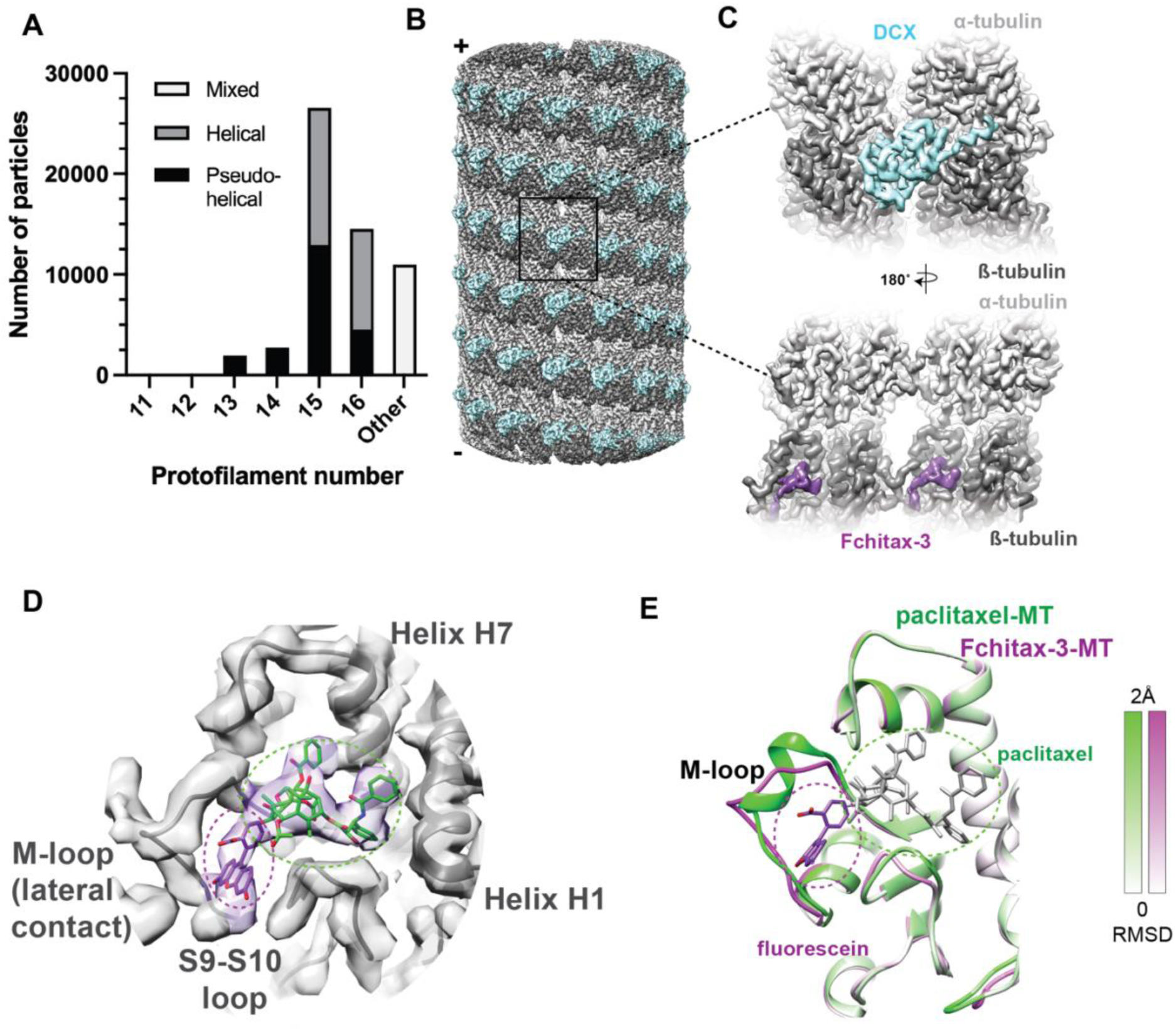
DCX binds 15-protofilament Fchitax-3 microtubules, lateral contacts of which are perturbed by the Fchitax-3 fluorophore. A) Stacked bar chart showing the number of particles from the cryo-EM data classified to the indicated architecture; B) Reconstruction of 15_4 microtubules bound with *a*-tubulin (silver), ß-tubulin (grey), and DCX (cyan), with plus and minus ends indicated; C) Asymmetric unit, comprising four tubulin monomers from adjacent protofilaments showing *a*-tubulin (silver), ß-tubulin (grey), DCX (cyan) and Fchitax-3 (purple), of the reconstruction viewed externally (top) and from the lumen (bottom); D) Density and model of ß-tubulin with Fchitax-3 density, paclitaxel model from PDB: 8UTO in the canonical pocket (green sticks) and fluorescein model (purple sticks) separately docked; E) Overlay of paclitaxel-bound ß-tubulin models with (purple scale) and without (green scale) fluorescein moiety attached, shaded by relative RMSD (PDB: 8UTO). Same view as D); Fchitax-3 is shown in stick representation (paclitaxel, green; fluorescein, purple).

Before assessing the effect of Fchitax-3 on the microtubule lattice, we confirmed that density corresponding to GTP and GDP is bound to *a*- and β-tubulin, respectively, consistent with hydrolysis of β-tubulin-bound GTP during microtubule assembly (Fig. S4A). One important structural change within microtubules on GTP hydrolysis is the transition from a so-called expanded to a compacted lattice, in which the overall length of the lattice-based tubulin dimer shrinks by ∼2% from >83 Å to <82 Å^28^. The tubulin lattice spacing in both our reconstructions was very similar: intra-dimer distance: 15_4: 41.85 ± 0.05 Å (n=3), 15_3: 42.05 ± 0.05 Å (n=4); inter-dimer distance: 15_4: 41.6 Å (n=3), 15_3: 41.4 Å (n=4), giving an overall dimer length of 83.5 Å in both reconstructions. Thus, although GDP is bound to β-tubulin, Fchitax-3 promotes an expanded lattice in these microtubules, in agreement with most previous reports of taxane stabilization^29–31^.

Consistent with this, well-defined density for Fchitax-3 was observed in the β-tubulin taxane pocket, allowing analysis of its binding mode and local structural changes in tubulin (Fig. 2C, purple density). The density corresponding to its paclitaxel moiety^3^ agrees well with previously determined structures^3,32^ (Fig. S3E, F, PDB:8UTO). The remaining Fchitax-3 density, which we interpret as the fluorescein fluorophore, projects from the taxane binding pocket towards the inter-protofilament lateral contact (Fig. 2D). This is consistent with the C7 position of fluorophore conjugation to paclitaxel^33^, and with antibody binding data indicating proximity of the fluorophore to the regular structural perforations between protofilaments^34^. In addition, our structure shows that the Fchitax-3 fluorophore contacts the base of the β-tubulin M-loop (Fig. 2D). Comparison of our 15_4 Fchitax-3-bound tubulin structure with a 15_4 paclitaxel-bound structure shows that the fluorophore of Fchitax-3 locally perturbs the M-loop structure and disrupts its short *a*-helix^32^ (Fig. 2E). Thus, although Fchitax-3 stabilizes microtubules analogously to other taxanes by occupying the canonical position within the β-tubulin lumenal pocket, its fluorescent moiety appears to act as a structural wedge at β-tubulin lateral contacts. If present during polymerization, this would narrow the inter-protofilament angle, thereby favouring an increase in the average protofilament number per microtubule. Taken together, our reconstructions show that Fchitax-3 stabilizes higher-protofilament-number microtubules and that these microtubules have expanded lattices.

### The N-DC domain of DCX is sufficiently flexible to bind between protofilaments in 15-protofilament microtubules

DCX is consistently observed to preferentially form 13-protofilament microtubules and bind compacted lattices^15,17,18,22^, so it was unexpected to observe it binding to 15-protofilament, expanded Fchitax-3 microtubules. We thus set out to define the structural basis of this interaction. Our 3D reconstruction protocol averages 8 nm spaced asymmetric units, yielding density corresponding to a single DC domain at the interdimer inter-protofilament interface^14,15^. In principle, this DC density could correspond to either N-DC, C-DC, or a mixture of both. Although the resolution of the density was not sufficient to visualize all amino acids, density corresponding to the longer (by 4 residues) loop3 of N-DC was prominent in the reconstruction (Fig. 3A-C). Further, there was no evidence of distinct sub-populations of N-DC and C-DC from 3D-classification of the DC domain density (Fig. S4B). Although the presence of the loop3 density does not exclude that C-DC is also bound in a subset of averaged asymmetric units (Fig. 3C), this density feature suggests that N-DC is the main binding element on Fchitax-3 microtubules. Furthermore, the prominent density N-terminal to the DC domain is more consistent with previous reconstructions of microtubule-bound N-DC than C-DC structures, in which this feature is not observed^15^ (Fig. 3B, C, right panels).

**Figure 3.**
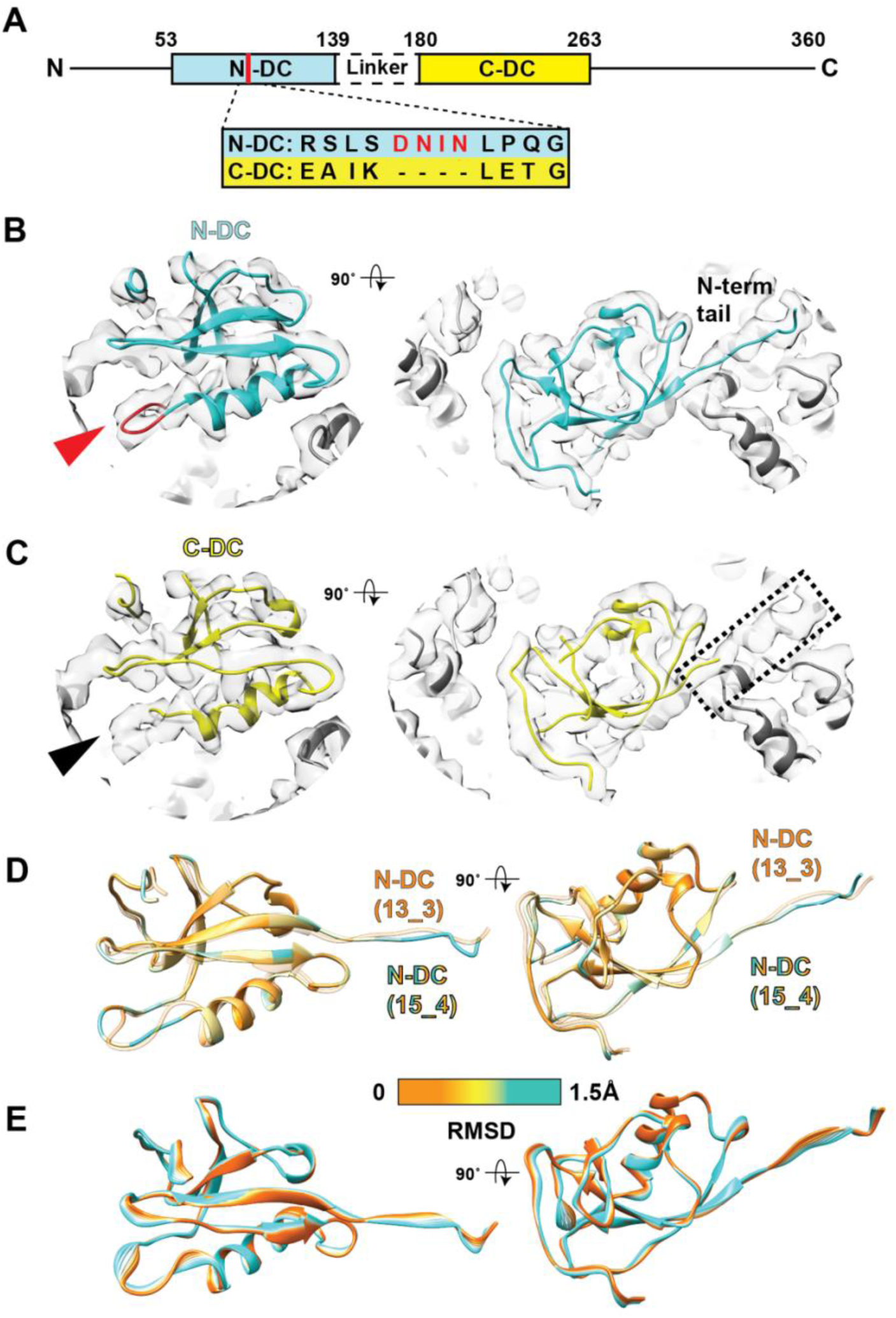
DCX binds to Fchitax-3-stabilized microtubules via the N-DC domain, which is flexibly accommodates in the smaller inter-protofilament binding space. A) Domain architecture of human DCX with the position of differentiating loop 3 sequences from N-DC and C-DC indicated. B) Density and model of 15_4 asymmetric unit showing, left DCX N-DC (cyan) at the inter-protofilament, inter-dimer binding site, viewed from the side, with N-DC loop3 highlighted in red and corresponding density indicated by red arrowhead; *a*-(silver) and ß-tubulin (dark grey); right N-DC viewed from the outside of the microtubule, with density corresponding to the characteristic N-DC N-terminal region indicated. C) Density and model of 15_4 asymmetric unit showing, left DCX C-DC (yellow, PDB:6RF2) at the inter-protofilament, inter-dimer binding site, viewed from the side; while the DC core is well accommodated by the density, density close to the shorter CDC loop3, indicated by black arrowhead, is unaccounted for; *a*-(silver) and ß-tubulin (dark grey); right C-DC viewed from the outside of the microtubule, with density corresponding to the characteristic N-DC N-terminal region unassigned when C-DC is docked. D) Overlay of N-DC domain of 15_4 microtubule (colored) and N-DC from 13_3 microtubule^15^ (orange) colored based on RMSD (scalebar provided), showing flexibility in the whole DC domain to accommodate the smaller space between protofilaments. View angles as in panels B, C. E) Overlay of 8 frames from a morph between N-DC from 13_3 microtubule (orange) and 15_4 microtubule (cyan) to highlight the size and direction of the changes undergone by N-DC to accommodate in different microtubules architectures.

Given that the angle between protofilaments is ∼3 ° narrower in 15-compared to 13-protofilament microtubules, we explored how N-DC is accommodated in the narrower inter-protofilament space. Comparison of the previously determined N-DC structure from a 13_3 microtubule with our 15_4 N-DC model shows that the domain flexes via slight repositioning of the loops throughout the DC domain (Fig. 3D, E). This behavior is consistent with previous evidence of domain plasticity, including loop position variability in N-DC structures from DCX and DCLK^35,36^ and with the structure of N-DC bound to 14-protofilament microtubules^15^.

### DCX tunes the expanded Fchitax-3 lattice into a conformation that favours binding

The flexibility of the N-DC domain described above explains how DCX binding can be accommodated within the narrower inter-protofilament valley of 15-protofilament microtubules. However, it does not account for DCX binding to the expanded lattice of Fchitax-3 microtubules. We therefore explored secondary-structure features within tubulin that undergo characteristic, coordinated conformational changes during the transition between lattice expansion and compaction. In particular, DCX binds close to both *a*-tubulin H8 helix, which undergoes a conformation adjustment, as does the so-called compaction switch of β-tubulin T5 loop, which is closed (expanded lattice) or open (compacted lattice)^31,37^ (Fig 4A-C). Strikingly, we found that despite the DCX-Fchitax-3 microtubules being longitudinally expanded, the structural signatures were more consistent with a compacted configuration – that is, DCX binding locally induces a hybrid lattice conformation that favours its binding despite being longitudinally expanded.

**Figure 4.**
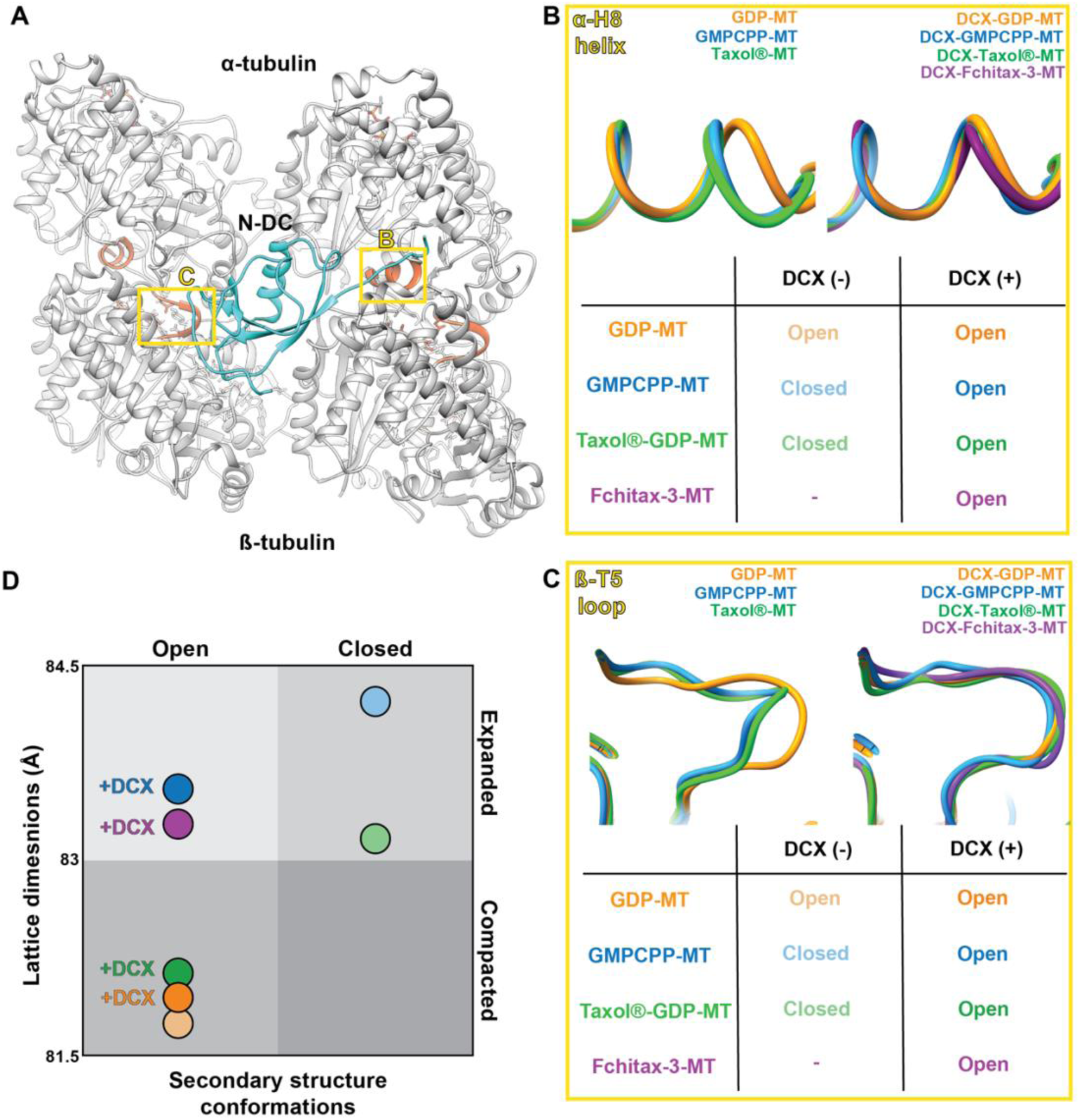
DCX tunes the expanded Fchitax-3 lattice into a binding-competent hybrid conformation. A) DCX binding site model viewed towards the microtubule surface, showing subunits from four tubulin dimers from adjacent protofilaments, *a*-tubulin (silver), ß-tubulin (grey), N-DC of DCX (cyan); two features within tubulin that undergo characteristic conformational changes between lattice expansion and compaction, *a*-tubulin H8 helix and β-tubulin T5 loop, are highlighted adjacent to the DCX binding site (red in the model, yellow rectangles); B) Comparison of conformational states observed in *a*-tubulin H8 helix in a set of microtubule reconstructions in the presence and absence of DCX (GDP-MT PDB:6DPV; GMPCPP-MT PDB:6DPU; Taxol®-GDP-MT PDB:6WVL; DCX-GDP-MT PDB:6REV; DCX-GMPCPP-MT PDB:6EVW; DCX-Taxol®-GDP-MT PDB:6EW0); C) Comparison of conformational states observed in β-tubulin T5 loop in a set of microtubule reconstructions in the presence and absence of DCX; PDBs as in B); D) Plot of lattice expansion/compaction vs *a*-tubulin H8 helix and of β-tubulin T5 loop structural states in the presence and absence of DCX. DCX binding promotes an open configuration of *a*-tubulin H8 helix and of β-tubulin T5 loop as it binds independently of longitudinal expansion/compaction.

To further explore this trend, we plotted these structural features from previously published reconstructions (by our group and others), in the presence and absence of DCX (Fig. 4D). Notably, apart from the structures in the current work, all DCX reconstructions were determined using microtubules copolymerized with DCX. Almost all the DCX reconstructions have longitudinally compacted lattices with structural features consistent with compacted conformations and with DCX’s binding preferences. Intriguingly, the one exception is the DCX-copolymerized GMPCPP microtubule structure, which has a longitudinally expanded lattice but compacted structural features (Fig. 4D, dark blue circle). These microtubules are of particular interest since the DCX-bound status of the GMPCPP microtubules appear to contradict the clearly poor binding of DCX to GMPCPP seeds (Fig. 1). However, we interpret this difference as being due to DCX having adapted these microtubules during polymerization to enable DCX binding, thereby producing a microtubule lattice with hybrid features, similar to those produced when DCX binds Fchitax-3 microtubules.

## DISCUSSION

Beyond their canonical structural and transport functions, microtubules are increasingly recognized as signalling hubs that regulate gene expression, cell behaviour and tissue-level processes^38,39^. Understanding how microtubule-binding proteins respond to changes in the microtubule lattice is therefore central to elucidating the mechanisms by which these signals are sensed and propagated. In this context, the mechanistic basis of how microtubule binding proteins respond to MSAs – as recently illustrated by RhoA signalling activation by taxanes^25^ - remains an important but complex facet of cytoskeleton biology. In our study, we used TIRF microscopy to explore binding by the neuronal protein DCX to a range of taxane-stabilized and taxane-seeded microtubules. In spite of its known preference for nucleating 13-protofilament microtubules with compacted lattices^17,18^, we found that DCX did not discriminate different MSA-stabilized microtubule populations even though taxanes promote lattice expansion and favor microtubules with between 12 and 15 protofilaments. The structural exploration of the interplay between DCX and Fchitax-3 binding prompted by these observations revealed a surprising degree of plasticity in the microtubule lattice.

We found that the fluorescein moiety of Fchitax-3 inserts at lateral contacts in 15-protofilament DCX-bound microtubules. This direct interaction of the fluorophore with β-tubulin is consistent with the dampening of its fluorescence on microtubule binding^40^. Insertion of the Fchitax-3 fluorophore at the ß-tubulin lateral contact appears to act as a wedge that narrows the inter-protofilament angle during polymerization and increases the average protofilament number in the resulting polymers. The perturbation of β-tubulin lateral contacts also points to a mechanism by which Fchitax-3 perturbs microtubule dynamics while still supporting polymerization and stabilization^8,9^. Although the taxane pocket is most accessible in lattice-incorporated tubulin^5^, the intermediate structural states through which tubulin passes as it polymerizes may also allow taxane binding^41^. The sporadic formation of lattice defects in microtubules grown in the presence of Fchitax-3 is thereby explained as stabilization of non-tubular polymers promoted by the fluorophore at the lateral contacts. Taken together, while fluorescent taxanes remain powerful tools for studying microtubules, our findings demonstrate that fluorophore identity and position is not a passive feature of fluorescent taxanes but an important determinant of their effects on microtubule lattice structure and dynamics.

As well as exhibiting variable protofilament number, microtubules undergo a longitudinal structural transition from an expanded, presumed stabilizing, GTP state to compacted, presumed destabilizing, GDP state^22,42^. It is commonly held that these states are binary in dynamic microtubules and that taxanes and other MSAs are stabilizing because they cause lattice expansion independent of nucleotide state. Intriguingly, however, DCX binding at the corner of four dimers imposes a hybrid lattice state on Fchitax-3 microtubules in which, despite longitudinal expansion, DCX moulds tubulin secondary structural elements towards conformations more consistent with compaction, DCX’s preferred lattice state. The saturated binding of DCX used in our cryo-EM experiments presumably facilitates this; in the TIRF experiments, however, the local concentration of DCX is lower and more variable. This explains the observed patchy binding behavior, likely due to the formation of structurally distinct regions with different DCX affinities within the same microtubule. Apparent discrepancies in the literature regarding DCX-taxane competition likely reflect differences in experimental conditions, including protein concentration, buffer composition, order of addition, and cellular context^21,22,43^. Our observations offer a framework through which these varied findings can be interpreted.

DCX’s ability to mould taxane-stabilized microtubule lattices depends on tubulin’s structural flexibility at its binding site. This implies that lattice expansion and compaction are not strictly binary and provide a mechanism for understanding why DCX binding does not straightforwardly report the lattice state. However, our analysis suggests this structural flexibility – particularly that of β-tubulin T5 – is constrained when GMPCPP is bound to the E-site. This means that in preformed GMPCPP microtubules, the expanded lattice is un-mouldable, explaining DCX’s poor binding. It is striking that while DCX can mould its binding site despite the expanded state of the Fchitax-3 lattice, a previous reconstruction of DCX bound to paclitaxel-stabilized microtubules exhibited a compacted lattice^23^. We speculate that the 15-protofilament architecture and/or the presence of fluorescein at the β-tubulin lateral contact prevent DCX from compacting these microtubules, whereas in paclitaxel-stabilized 13-protofilament microtubules, full compaction is possible. In contrast, a recent survey of MSAs found that a set of fluorescent taxanes, including Fchitax-3, exhibited compacted lattices, whereas other taxanes promoted expansion^44^. This study also found, however, that these fluorescent taxane-bound microtubules were built primarily from 14 protofilaments, in contrast to our data presented here and that of the same group in a different study^9^. While it is not immediately obvious how to reconcile these observations, differences in polymerization conditions may again influence the specific properties of the resulting microtubules. Taken together, these findings suggest that the interplay between polymerization and stabilization conditions in determining microtubule lattice state and architecture is more complex than currently recognized.

Our findings show that DCX binding to microtubules is more adaptable than previously appreciated. Although DCX preferentially nucleates and binds 13-protofilament microtubules with a compacted lattice, and can exhibit cooperativity^14,17,18,20,22,23^, its inter-protofilament binding site was originally characterized on 14-protofilament paclitaxel-stabilized microtubules^17^. Together with numerous other reports of DCX binding to paclitaxel-stabilized microtubules^19,20,23,36^, our data now imply that earlier studies underestimated the breadth of DCX’s microtubule-binding capabilities^17^. We previously proposed that each DC domain in DCX has distinct roles, with C-DC promoting nucleation of the characteristic 13 protofilament architecture then handing over to N-DC – with its larger binding footprint – for lattice stabilization^15^. Consistent with this, we found that N-DC flexibly adapts to the smaller binding site found in 15-protofilament lattice.

Although DCX is essential for neuronal migration during human brain development^10,11^, the underlying mechanisms remain incompletely understood. While DCX has been implicated in microtubule bundling, resistance to migration-associated forces, and regulation of tubulin post-translational modification^45–47^, microtubules in DCX-knockout neurons retain their canonical 13-protofilament architecture, arguing against a primary role in architecture specification^48^. Instead, the increased lattice discontinuities observed in the absence of DCX suggest a role in maintaining microtubule integrity. The 15-protofilament Fchitax-3 microtubules analysed here appeared free of lattice defects, despite reports that Fchitax-3 can induce ruptures^8^, consistent with DCX contributing to resistance to, or repair of, lattice damage during migration^46,48^. Given the growing clinical importance of tubulin-targeting agents^49,50^, understanding how such compounds interact with neuronal microtubule-associated proteins remains an important goal. DCX therefore provides a valuable readout of microtubule lattice allostery.

## METHODS

### In vitro microtubule seed preparation using GMPCPP and MSAs

Stabilized microtubule seeds for in vitro assays were assembled as described previously^8,9^. Briefly, microtubule seeds stabilized with GMPCPP were prepared by performing microtubule polymerization and depolymerization cycles twice in the presence of GMPCPP. A solution of porcine brain tubulin containing 12% HiLyte 488^TM^ labelled tubulin and 18% biotin-labelled tubulin was polymerized in MRB80 buffer (80 mM K-PIPES, pH 6.8, 4 mM MgCl_2_, 1 mM EGTA) in the presence of 1 mM GMPCPP. After polymerization, centrifugation was performed in an airfuge for 5 min at 119,000g. The pellet was resuspended in MRB80 buffer, then depolymerized on ice for 20 min. Repolymerization of the solution was performed in the presence of more GMPCPP. After the second polymerization and pelleting, the resulting GMPCPP stabilized microtubule seeds were stored in MRB80 buffer supplemented with 10% glycerol.

For the preparation of various MSA-stabilized microtubule seeds (paclitaxel, docetaxel and Fchitax-3), porcine brain tubulin solution, containing HiLyte 488^TM^ labelled tubulin and biotin-labelled tubulin was incubated in MRB80 buffer at 37°C for 20 min in the presence of 1 mM GTP and 20 μM of each MSA. After 20 min, polymerized tubulin solution was diluted 5 times with pre-warmed 20 µM MSAs and incubated further for 5 min. After incubation, centrifugation was performed at 17000 g for 15 min at 30°C.

The obtained pellet was resuspended in MRB80 buffer containing 20 μM of the respective MSAs and stored at RT with light protection.

### Tagged Doublecortin purification

N-terminal-TEV bio-tagged mCherry-Doublecortin (DCX, mouse variant) cloned in PTT5 vector was purified using the protocol as described previously^51^. Briefly, Bio-TEV-mCherry-DCX was purified from transiently transfected HEK-293T cells using streptavidin affinity purification followed by TEV protease-mediated elution. Cells were cultured in 50% DMEM with 50% Ham’s F-10 supplemented with 10% FBS and were co-transfected with Bio-TEV-mCherry-DCX and BirA expression plasmids using polyethyleneimine (PEI). After 24 h, cells were harvested, washed with ice-cold PBS, and lysed in lysis buffer (50 mM HEPES, pH 7.4, 300 mM NaCl, 1 mM MgCl₂, 0.5% Triton X-100, 1 mM DTT, and 1X protease inhibitor cocktail). Following clarification by centrifugation (17000g, 15 min, 4°C), the lysate was incubated with chicken egg albumin-blocked Dynabeads M-280 Streptavidin (Invitrogen) for 45 min at 4°C. Beads were washed three times with wash buffer (50 mM HEPES, pH 7.4, 300 mM NaCl, 1 mM MgCl₂, 0.5% Triton X-100, and 1 mM DTT), followed by three washes with TEV cleavage buffer (50 mM HEPES, pH 7.4, 150 mM NaCl, 1 mM MgCl₂, 1 mM EGTA, 0.05% Triton X-100, and 1 mM DTT). Bound Bio-TEV-mCherry-DCX was released by incubation with recombinant TEV protease (Sigma-Aldrich) in TEV cleavage buffer for 2 h at 4°C. The eluate was snap-frozen in liquid nitrogen and stored at −80°C until use for in vitro microtubule assays.

### In vitro assay for microtubule-doublecortin interaction using TIRF microscopy

In vitro chamber preparation, microtubule seed imaging, microtubule dynamics assay, and subsequent image acquisition by TIRF microscopy were performed as described in detail previously^8,9^. In brief, in vitro TIRF microscopy imaging flow chambers were prepared with two parallel strips of double-sided tape on microscopic slides with plasma-cleaned glass coverslips. Flow chambers were first incubated with 0.2 mg/ml PLL-PEG-biotin and then 1 mg/ml NeutrAvidin in MRB80 buffer. Flow chambers were further incubated with microtubule seeds as indicated, followed by treatment with 1 mg/ml κ-casein. The reaction mixtures of 15 μM porcine brain tubulin containing 0.5 µM HiLyte 488^TM^ labelled tubulin, supplemented with 50 nM or 200 nM DCX (as indicated), 20 nM GFP-EB3 (as indicated), 0.1% methylcellulose, 0.2 mg/ml κ-casein, 50 mM KCl, 1 mM GTP and oxygen scavenger mixture (50 mM glucose, 200 μg/ml catalase, 4 mM DTT and 400 μg/m glucose oxidase in MRB80 buffer) were centrifuged in an airfuge at 119,000g for 5 min and were added to the flow chambers. Vacuum grease was used for closing flow chambers and TIRF microscopy was performed. Samples were imaged at 30°C.

### Image acquisition with TIRF microscopy

All in vitro reconstitution assay samples were imaged using a TIRF microscope setup (inverted research microscope, Nikon Eclipse Ti-E). The system has an automated perfect-focus module (Nikon) in conjunction with a Nikon CFI Apo TIRF ×100 objective lens (numerical aperture 1.49, oil-immersion). TIRF illumination was achieved via a motorized TIRF-E illuminator (customized by Roper Scientific/PICT-IBiSA Institut Curie). Dual-colour (red/Green) simultaneous fluorescence imaging was enabled with a triple-band TIRF polychroic mirror (ZT405/488/561rpc, Chroma) paired with a corresponding triple-band emission filter (ZET405/488/561 m, Chroma) mounted within the metal cube (Chroma, model 91032), Spectral separation of emission signals was further achieved using an Optosplit III image splitter (Cairn Research), fitted with a dual-emission filter set comprising ET525/50 m and ET630/75 m filters alongside a T585LPXR dichroic mirror (Chroma). The sample temperature was maintained using a stage-top incubation chamber (INUBG2E-ZILCS, Tokai Hit). Images were captured with either a Photometrics Evolve 512 EMCCD camera or a Photometrics CoolSNAP HQ2 CCD camera (Both Roper Scientific), with acquisition managed through MetaMorph 7.7 software (Molecular Devices).

### Analysis of DCX-decorated region relative to microtubule length

The extent of DCX lattice occupancy was quantified using the Fiji plugin (Curve trace 0.3.5; https://github.com/UU-cellbiology/CurveTrace). Microtubules were detected and traced in the green fluorescence channels using Steger’s line detection algorithm and the corresponding mCherry-DCX (red channel) binding along the lattice was measured for microtubules stabilized with paclitaxel, GMPCPP, docetaxel, or Fchitax-3. Relative DCX binding (%) across all microtubule seeds was calculated and plotted using GraphPad Prism software. Further, to assess the spatial distribution of DCX along individual microtubules, fluorescence intensity line scans were generated using the Plot Profile function in Fiji/ImageJ across representative microtubules stabilized with paclitaxel, GMPCPP, docetaxel, or Fchitax-3. The resulting fluorescence intensity profiles were analyzed and plotted using GraphPad Prism.

### Cryo-EM sample preparation

The full length human doublecortin sequence was subcloned into the pNic28-Bsa4 vector (Structural Genomics Consortium, Oxford, UK), with an N-terminal TEV protease-cleavable His-tag. It was expressed in One Shot™ BL21 Star™ (DE3) Chemically Competent *E. coli* cells (Thermo Fisher) and purified according to Manka & Moores^15^.

Fchitax-3 stabilized microtubules were prepared as above in the Akhmanova lab and sent to the Moores lab for cryo-EM. Fchitax-3 microtubules (20 μM Fchitax-3 per 15 μM tubulin) were diluted to 3 μM tubulin in BRB80 buffer (80 mM PIPES [piperazine-N,N′-bis(2-ethanesulfonic acid)] pH 6.8, 1 mM EGTA [ethylene glycol-bis(β-aminoethyl ether)-N,N,N’,N’-tetraacetic acid], 1 mM MgCl2, 1 mM DTT [dithiotreitol]) containing 1 mM GTP. C-flat 2/2 Holey Carbon (Electron Microscopy Sciences) or Lacey (Agar) grids were glow-discharged in air using a Harrick Plasma Cleaner at 0.2-0.3 Torr for 40 seconds. 4 μl of diluted Fchitax-3-microtubules were applied to grids, grids were briefly blotted on the side with filter paper, 4 μl of 50 μM doublecortin (DCX) solution in BRB80 buffer was added, and the grids were transferred into the chamber of the Vitrobot Mark IV (Thermo Fisher Scientific). The grids were incubated in the Vitrobot chamber for 1 min at 30 °C and 95 % humidity, before blotting and plunge freezing in liquid ethane.

### Cryo-EM data collection

Data were collected on a Titan Krios D3771 microscope (Thermo Fisher Scientific) operating at an accelerating voltage of 300 kV attached to a BioQuantum K2 direct electron detector (Gatan) and post-column GIF energy filter (Gatan) with a slit width of 20eV. The samples were imaged using the automated EPU software. 2660 movies were collected with defocus ranging between -0.4 to -4.5 μm. 9-second exposures over 35 frames delivered a total electron dose of 26.25 e^-^/Å^2^ to the specimen per movie. The magnified pixel size was 1.38 Å. Data collection statistics are provided in Table 1.

**Table 1.**
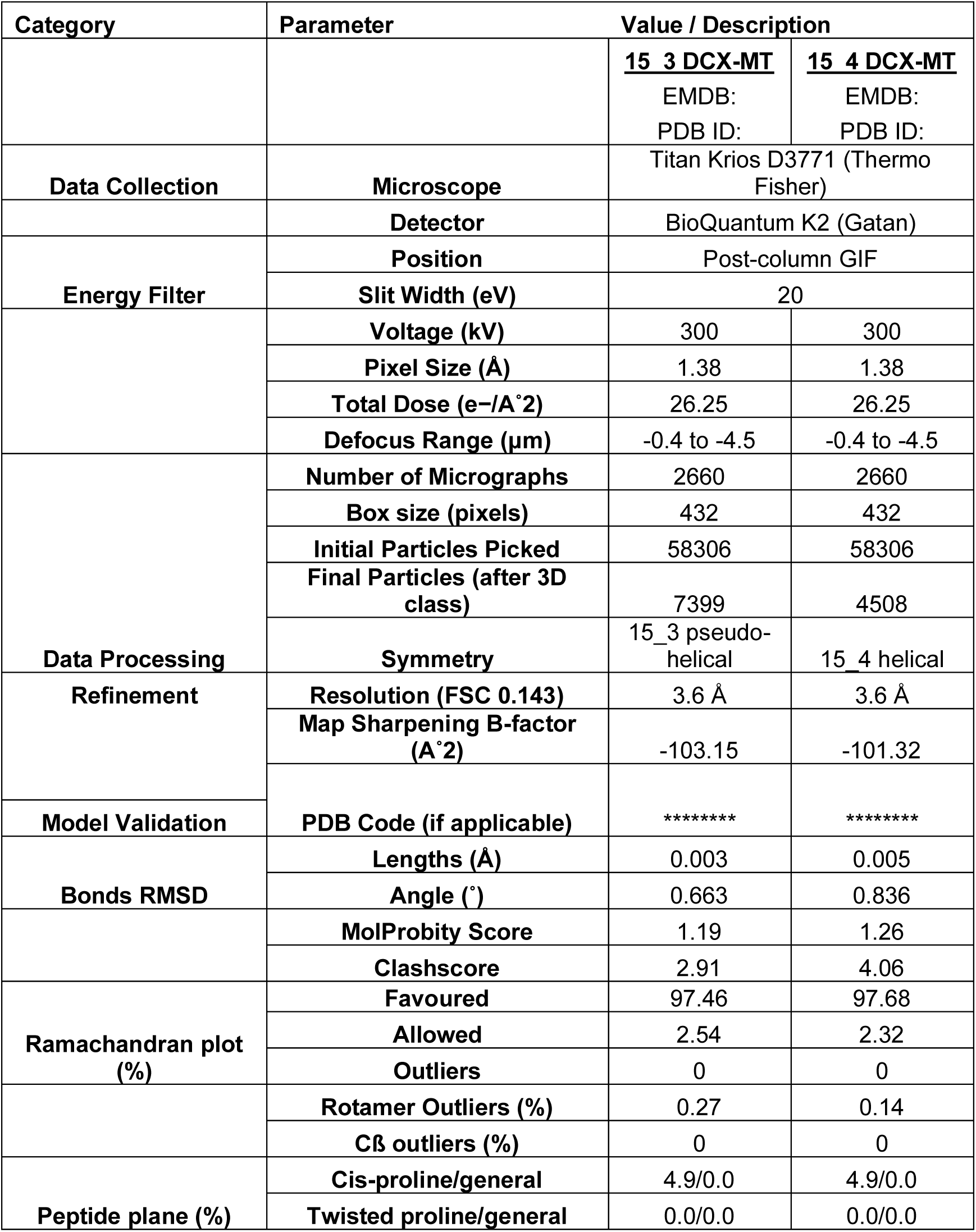

| Category | Parameter | Value / Description |  |
|---|---|---|---|
|  |  | 15_3 DCX-MT<br>EMDB:<br>PDB ID: | 15_4 DCX-MT<br>EMDB:<br>PDB ID: |
| Data Collection | Microscope | Titan Krios D3771 (Thermo Fisher) |  |
|  | Detector | BioQuantum K2 (Gatan) |  |
| Energy Filter | Position | Post-column GIF |  |
|  | Slit Width (eV) | 20 |  |
|  | Voltage (kV) | 300 | 300 |
|  | Pixel Size (Å) | 1.38 | 1.38 |
|  | Total Dose (e−/A˚2) | 26.25 | 26.25 |
|  | Defocus Range (µm) | -0.4 to -4.5 | -0.4 to -4.5 |
| Data Processing | Number of Micrographs | 2660 | 2660 |
|  | Box size (pixels) | 432 | 432 |
|  | Initial Particles Picked | 58306 | 58306 |
|  | Final Particles (after 3D class) | 7399 | 4508 |
|  | Symmetry | 15_3 pseudo-helical | 15_4 helical |
| Refinement | Resolution (FSC 0.143) | 3.6 Å | 3.6 Å |
|  | Map Sharpening B-factor (A˚2) | -103.15 | -101.32 |
| Model Validation | PDB Code (if applicable) | ******** | ******** |
| Bonds RMSD | Lengths (Å) | 0.003 | 0.005 |
|  | Angle (˚) | 0.663 | 0.836 |
|  | MolProbity Score | 1.19 | 1.26 |
|  | Clashscore | 2.91 | 4.06 |
| Ramachandran plot (%) | Favoured | 97.46 | 97.68 |
|  | Allowed | 2.54 | 2.32 |
|  | Outliers | 0 | 0 |
|  | Rotamer Outliers (%) | 0.27 | 0.14 |
|  | Cß outliers (%) | 0 | 0 |
| Peptide plane (%) | Cis-proline/general | 4.9/0.0 | 4.9/0.0 |
|  | Twisted proline/general | 0.0/0.0 | 0.0/0.0 |

### Cryo-EM image processing

Fchitax-3-DCX-microtubule datasets were processed in RELION v3.0 and v3.1^52,53^. In RELION v3.0, movie frames were aligned, summed, and dose-weighted using MotionCorr v2.0^54^, generating micrographs at 1.38 Å/pixel. CTF parameters were estimated with CTFFind v4.1.14^55^. Microtubule coordinates were manually picked in helix mode and extracted as helices (110 Å inner, 390 Å outer diameter) with 4× binning (5.52 Å/pixel) and 82 Å inter-box distance. Extracted microtubule segments were further processed in RELION v3.1 using a customized MiRP v2.0 workflow^56^.

To determine the distribution of protofilament architectures in the extracted data, the particles were subjected to iterative 3D and 2D classifications as follows. Microtubule segments were averaged into super-particles, which were aligned and classified in 3D using six undecorated microtubule references (11–16 protofilaments) generated in UCSF Chimera v1.18^57^. The references were low-pass filtered to 40 Å, while the particles were aligned using Fourier components up to a resolution of 20 Å (E-step resolution limit). The alignment used 1.8° angular sampling intervals and a 10-pixel translational range in 1-pixel steps. Classified segments from each microtubule were then assigned to their modal protofilament class using the protofilament sorting algorithm in MiRPv2.0^58^. To further resolve any remaining heterogeneity in each protofilament architecture, the sorted particles were further classified in 2D without re-alignment and the diameter of the microtubules in the classes was measured. Particles belonging to classes that showed the same diameter were manually grouped into the same protofilament class. Particles belonging to the 15-protofilament architecture were then segregated in 3D based on helical start-number into pseudo-helical (3-start) and helical (4-start) architectures. This was performed using two references of DCX-decorated microtubules corresponding to 3-start and 4-start architectures respectively, generated in UCSF Chimera.

For the 15_3 microtubules, the particles were aligned to a common register to locate the point of asymmetry i.e. the seam. This was achieved by refining the rotation angle (ø) and translational shifts (XY) through using MiRP v2.0^58^. This was followed by a final alignment-free 3D classification using references with 15 different seam positions, each corresponding to a different protofilament, to filter out any poorly aligned particles. The aligned particles were then extracted at original pixel size and used to refine the corresponding seam-reference without the application of symmetry. The sampling parameters for alignment were limited to local searches of 2° and 5-pixels in 0.9° and 0.5-pixel increments respectively. The refined particles were symmetry-expanded across all 15 protofilaments and the Rot-(ø)-angle priors modified using helical parameters that were determined from the refined map using RELION’s Helix Toolbox. The expanded particles were used to further refine the map, which was sharpened using RELION’s LocalRes tool^53,59^. The presence of a seam resulted in the αß-tubulin register being lost in all but one of the protofilaments across from the seam. Thus, the asymmetric unit for model building was extracted from the two protofilaments across from the seam in the sharpened map. It comprises a single DCX domain sandwiched between two adjacent tubulin dimers.

For the symmetric 15_4 microtubule, the particles were aligned in a single step of refinement using the 15_4 decorated microtubule reference without the application of symmetry. The resultant reconstruction was then refined with the application of helical symmetry, using parameters derived from RELION’s helix toolbox, and sharpened by RELION’s LocalRes^53,59^. The tetrameric asymmetric unit was extracted for model building.

### Model Building and Refinement

Initial models for N-terminal DC domain of *Homo sapiens* Doublecortin (UniProt: O43602) spanning residues 44-142 as well as *Sus scrofus* α1b-(UniProt: Q2XVP4) and ß1-tubulin (UniProt: P02554) were generated separately using AlphaFold v3.0^60^. Fchitax-3 was modelled as a combination of two ligands - paclitaxel (PDB: TA1) and fluorescein (PDB: FLU). For each asymmetric unit-15_3 and 15_4 microtubules, the models of the two tubulin dimers and N-DC were docked into the density and the ligands into their respective difference maps using UCSF Chimera’s Fit-In-Map tool and the residues that were not represented by the density were trimmed. The fit of the combined protein-ligand model was refined manually in ISOLDE^61^ followed by real space refinement in PHENIX v1.21^62^. The refinement strategy in PHENIX comprised of global geometry minimisation, Anisotropic Displacement Parameter (ADP) refinement, and local grid searches, with restraints for secondary structure, Ramachandran angles, and rotamers in the protein as well as restraints from the CIF dictionaries of the ligands. All refinement was performed on maps that were sharpened by the LocalRes tool in RELION v3.1.

To calculate inter- and intra-dimer distances in the microtubule lattice, each dataset was first split into four (15_3) and three (15_4) subsets, and each subset was used to independently calculate a 3D reconstruction. In each reconstruction, two copies of the tetrameric asymmetric unit model (2α tub-NDC-2ß tub) was fitted as a rigid body such that they occupy adjacent positions on the same pair of protofilaments. Lattice spacing was then measured as the distance between the ribose moiety of the nucleotides^63^, using The PyMOL Molecular Graphics System, Version 3.1.6 Schrödinger, LLC.

### Resource table

| Reagent or Resources | Source | Identifier |
|---|---|---|
| Paclitaxel | Enzo Life Sciences | Cat#T7402 |
| Docetaxel | Sanofi-Aventis | Aventis, Schiltigheim, France |
| Fchitax-3 | A kind gift from CIB Margarita Salas / Dr. Wei-Shuo Fang (Institute of Materia Medica, Beijing) | N/A |
| GMPCPP | Jena Bioscience | Cat#NU-405L |
| GTP | Sigma-Aldrich | Cat#G8877 |
| Triton X-100 | Sigma-Aldrich | Cat#93443 |
| PIPES (1,4-piperazinediethanesulfonic acid) | Sigma-Aldrich | Cat#80635 |
| Magnesium chloride | Sigma-Aldrich | Cat#M8266 |
| Potassium hydroxide | Sigma-Aldrich | Cat#221473 |
| Potassium chloride | Sigma-Aldrich | Cat#P9541 |
| Glucose | Sigma-Aldrich | Cat#G7021 |
| Albumin from chicken egg white | Sigma-Aldrich | Cat#A5503 |
| Glucose oxidase (from *Aspergillus niger*) | Sigma-Aldrich | Cat#G7141 |
| Catalase (from bovine liver) | Sigma-Aldrich | Cat#C9322 |
| Dithiothreitol (DTT) | Sigma-Aldrich | Cat#R0861 |
| κ-Casein | Sigma-Aldrich | Cat#C0406 |
| Albumin from Chicken Egg white | Sigma-Aldrich | Cat#A5503-10G |
| Methylcellulose | Sigma-Aldrich | Cat# M0512 |
| Ethylene glycol-bis(2-aminoethylether) N,N,N′,N′-tetraacetic acid (EGTA) | Sigma-Aldrich | Cat#E4378 |
| NeutrAvidin | Invitrogen | Cat#A-2666 |
| Dynal streptavidin beads M-280 | Invitrogen | Cat#112-06D |
| Dynal magnetic separator MPC-S | Invitrogen | Cat#A13346 |
| Complete Protease inhibitor | Roche | Cat#04693116001 |
| Tubulin (porcine) | Cytoskeleton | Cat#T240-C |
| HiLyte 488<sup>TM</sup> labelled tubulin | Cytoskeleton | Cat#TL488M |
| Biotin labelled tubulin | Cytoskeleton | Cat#T333P |
| Biotinylated poly(L-lysine)-*g*-poly(ethylene glycol) (PLL-PEG-biotin) | Surface Solutions | PLL(20)-g[3.5]-PEG(2)/PEG(3.4)-biotin(50%) |
| C-Flat™ Holey Carbon Grid 2.0 µm hole 2.0 µm space 300 Mesh | Electron Microscopy Sciences | CF-223C-25 |
| **Software and Algorithms** |  |  |
| ImageJ | NIH | https://imagej.nih.gov/ij/ |
| Metamorph | MolecularDevices | https://www.moleculardevices.com/products/cellular-imaging-systems/acquisition-and-analysis-software/metamorph-microscopy |
| GraphPadPrism | GraphPad | https://www.graphpad.com/scientific-software/prism/ |
| Curve trace 0.3.5 plugin | EugeneKatrukha | https://github.com/UU-cellbiology/CurveTrace |
| KymoResliceWide plugin | EugeneKatrukha | https://github.com/ekatrukha/KymoResliceWide |

## DATA AVAILABILITY

The cryo-EM reconstructions generated in this study have been deposited in the Electron Microscopy Data Bank under accession code EMD-****** (15_4) and EMD ****** (15_3). The corresponding structural models generated in this study have been deposited in the Worldwide Protein Data Bank under the accession codes PDB: ******** (15_4) and PDB: ******** (15_3).

## ACKNOWLEDGEMENTS

We thank former and current members of the Moores, Akhmanova, and Rai laboratories for critical discussion. Cryo-EM work was supported by the Medical Research Council, U.K. (MR/R00352/1 and MR/Y000633/1 to C.A.M.). Cryo-EM data were collected at the ISMB EM facility (Birkbeck College, University of London) with financial support from the Wellcome Trust (202679/Z/16/Z and 206166/Z/17/Z). We thank Natasha Lukoyanova and Shu Chen for electron microscope support, and David Houldershaw for computational support at Birkbeck. A.R. thanks DBT sponsored Ramalingaswami-re-entry fellowship grant and GSBTM grant (202425_33).

## Declaration of generative AI and AI-assisted technologies in the manuscript preparation process

During the preparation of this work, the authors used Microsoft Co-Pilot for grammar correction. The authors reviewed and edited the output as needed and take full responsibility for the content of the published article.

## SUPPLEMENTARY FIGURES

**Figure S1.**
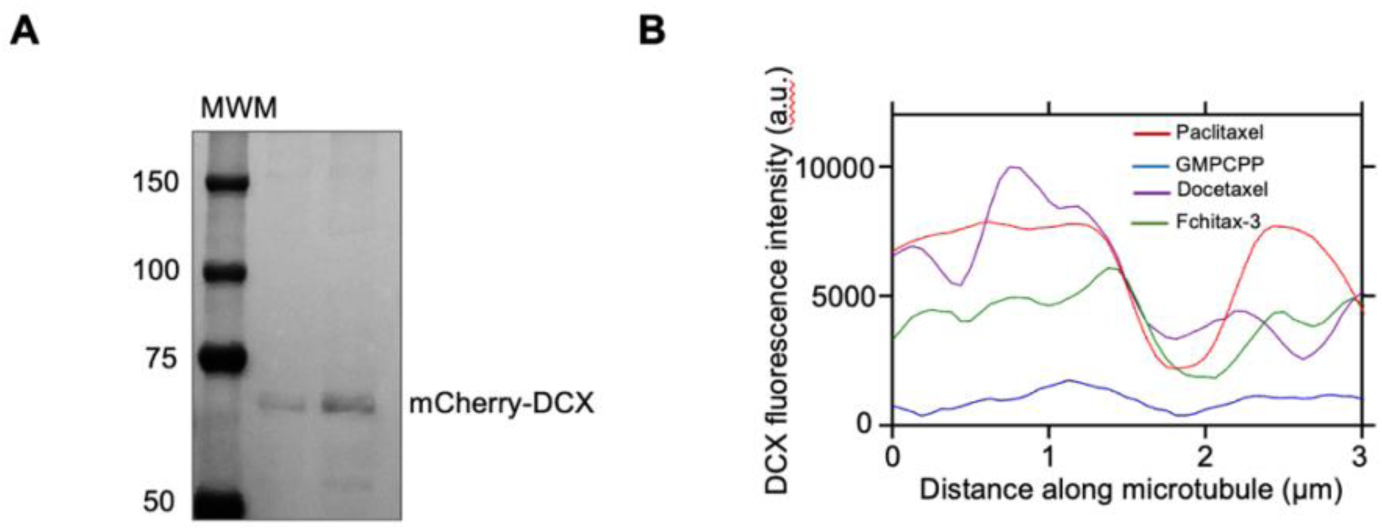
Characterization of mCherry-DCX and additional analysis of DCX binding. A) Coomassie blue stained gel showing purified TEV bio-tagged mCherry-Doublecortin (DCX) used for the TIRF in vitro reconstitution assays. Molecular weight markers (MWM) are indicated. B) Representative fluorescence intensity line scans of mCherry-DCX along individual microtubules. The line traces show that DCX is not uniformly distributed along the microtubule lattice but instead exhibits discrete regions of high and low fluorescence intensity, indicating discontinuous lattice occupancy.

**Figure S2:**
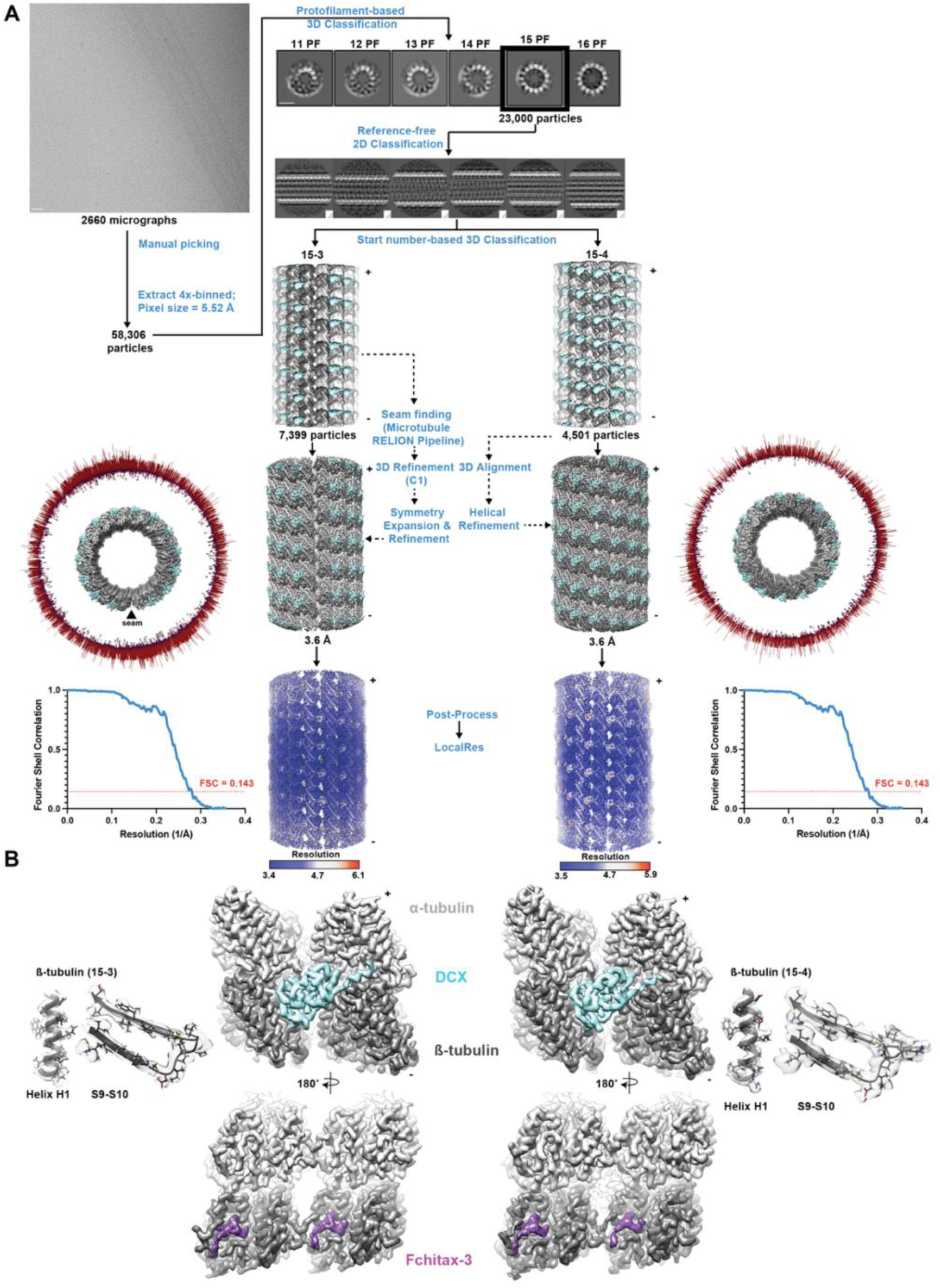
Cryo-EM reconstruction workflow of DCX-bound Fchitax-3 microtubules and exemplar reconstruction density quality. A) Cryo-EM reconstruction workflow. Steps of the workflow are indicated in blue. A representative micrograph is shown. Scale bar = 30 nm. Cross-sections of classes obtained from reference-based 3D classification are shown with corresponding protofilament (PF) architectures specified above. Scale bar = 15 nm. Subsequent reference-based 2D classification was performed on particles of the 15 PF class to eliminate remaining heterogeneity through visual inspection of class averages. EM reconstructions are shown in grey with microtubule plus/minus ends indicated. Angular distribution of particles used in 3D refinement are represented as cylindrical histogram plots around each microtubule (top view) and the seam is indicated (arrowhead). Corresponding FSC plots and density coloured by local resolution are also shown. B) Cryo-EM density of DCX-bound asymmetric unit segmented from each reconstruction, viewed from outside (top) and lumen (bottom). Density for the α-tubulin, β-tubulin, Fchitax-3 and DCX is shown in light, dark grey, purple and cyan respectively. In each case, representative regions of the reconstruction are shown with the fitted model (far left, far right, cartoon representation with side chains).

**Figure S3:**
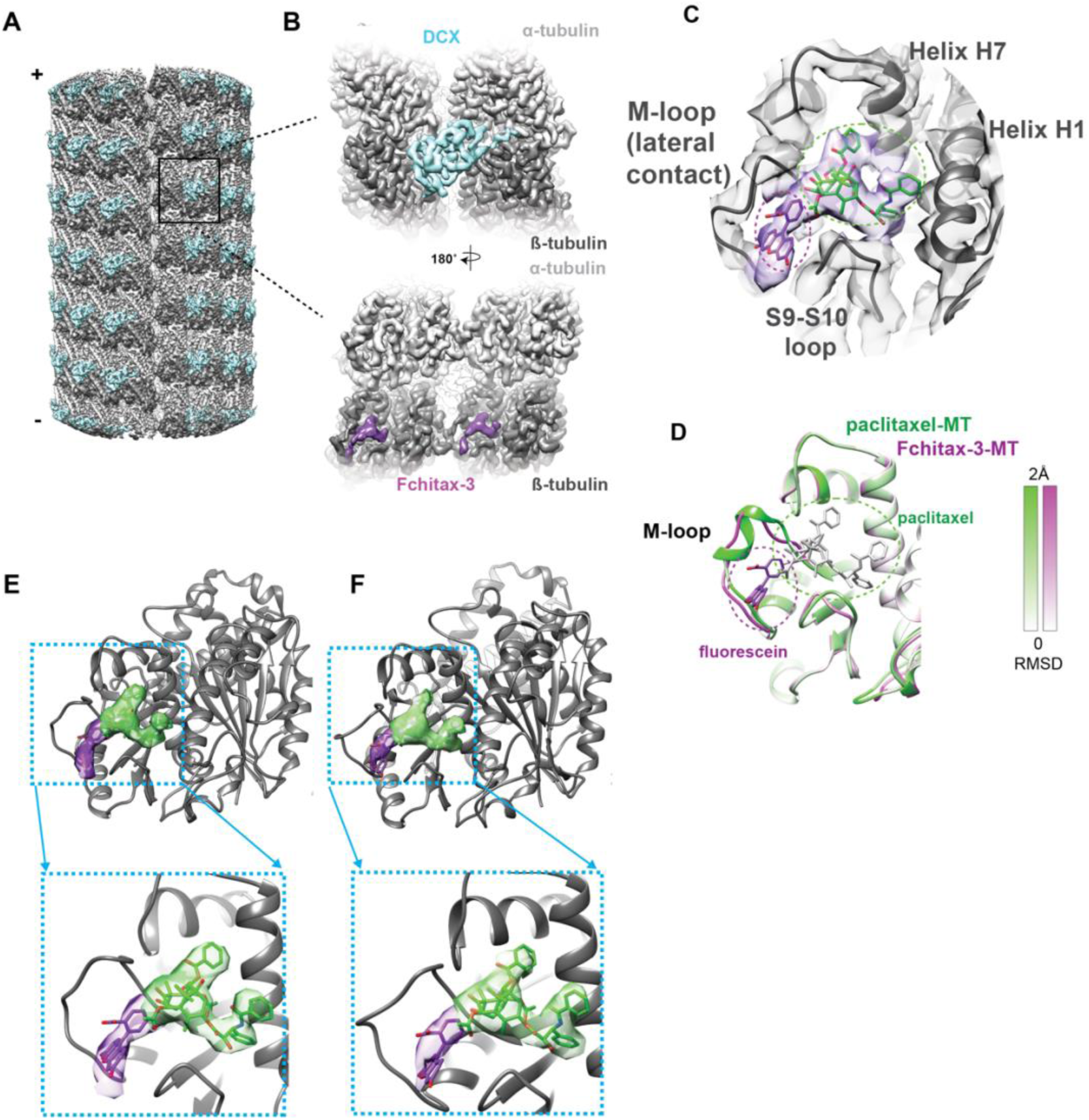
The mechanism by which doublecortin binds to Fchitax-3-stabilized 15_3 microtubules is the same as that of 15_4 microtubules, apart from at the seam. A) Reconstruction of 15_3 microtubules bound with *a*-tubulin (silver), ß-tubulin (grey), and DCX (cyan), with plus and minus ends indicated; B) Asymmetric unit, comprising two tubulin dimers from adjacent protofilaments showing *a*-tubulin (silver), ß-tubulin (grey), DCX (cyan), and Fchitax3 (purple). of the reconstruction viewed externally (top) and from the lumen (bottom); C) Density and model of ß-tubulin with Fchitax-3 density, paclitaxel model from PDB: 8UTO in the canonical pocket (green sticks) and fluorescein model (purple sticks), separately docked; D) Overlay of paclitaxel-bound ß-tubulin models with (purple scale) and without (green scale) fluorescein moiety attached, shaded by relative RMSD (PDB: 8UTO). Same view as C); Fchitax-3 is shown in stick representation (paclitaxel, green; fluorescein, purple). E) Density of Fchitax-3 in 15_4 microtubules bound to ß-tubulin (top) and an inset focusing on the ligand density (bottom); Fchitax-3 shown as a combination of fluorescein (purple) and paclitaxel (green), was obtained as a difference between the reconstructed density and the protein model-derived density; ß-tubulin is shown as grey ribbon-model. F) Density of Fchitax-3 in 15_3 microtubules bound to ß-tubulin (top) and an inset focusing on the ligand density (bottom); Fchitax-3 shown as a combination of fluorescein (purple) and paclitaxel (green) was obtained as a difference between the reconstructed density and the protein model-derived density; ß-tubulin is shown as grey ribbon-model.

**Figure S4.**
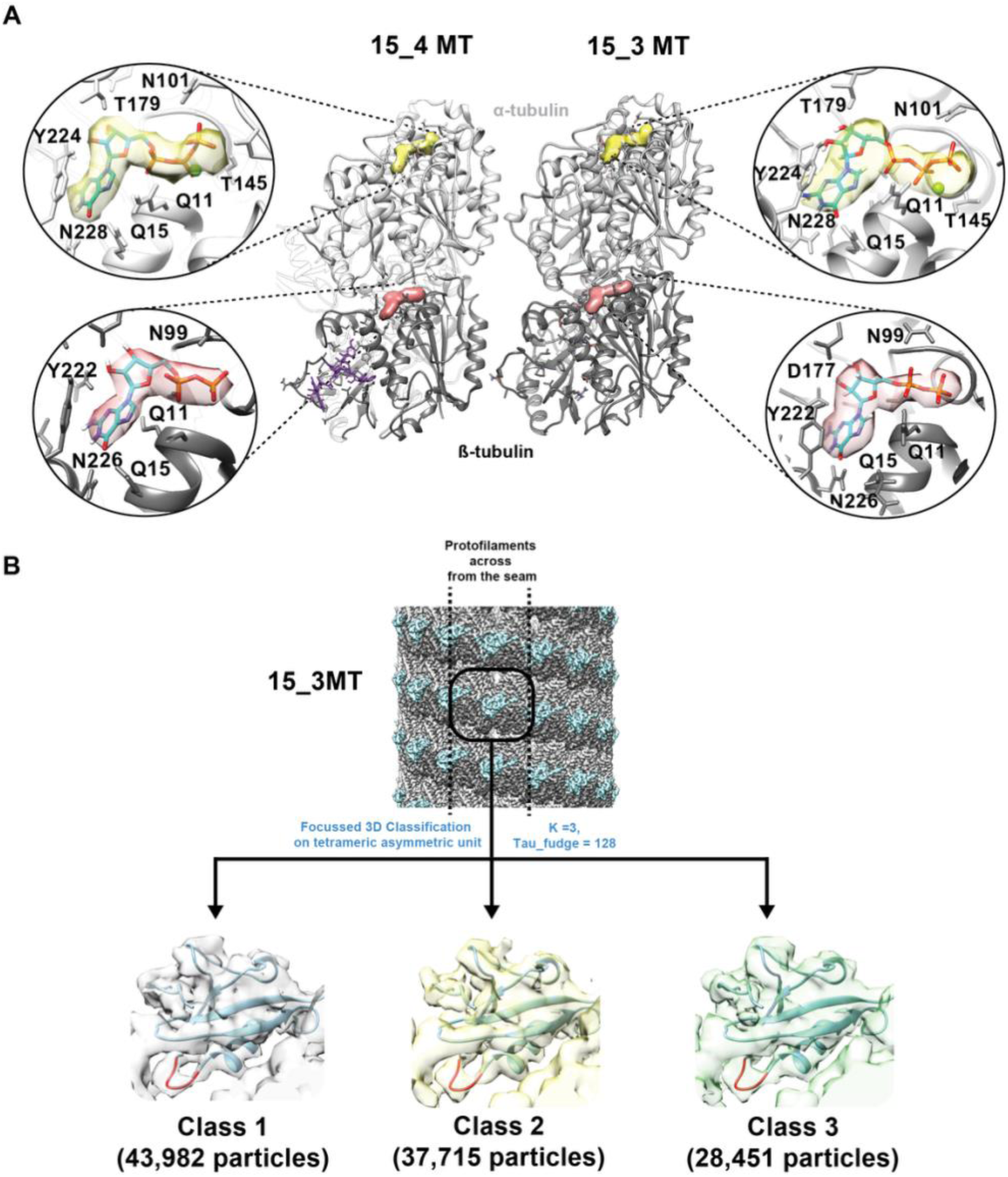
Cryo-EM nucleotide data quality control and DC domain focused classification. A) Nucleotide density bound to *a*- and ß-tubulin in 15_3 and 15_4 microtubule reconstructions. Each reconstruction is viewed from the lumenal side, tubulins are depicted as ribbon models with density corresponding to bound ligand calculated from the difference between reconstructed density and protein model-derived density. *a*-tubulin-bound GTP is shaded yellow and ß-tubulin-bound GDP shaded orange. ß-tubulin-bound Fchitax-3 is shown in purple stick representation. B) 3D classification of the DC domain in symmetry expanded 15_3 data produces three classes which all show clear density corresponding to N-DC loop3.

